# Neural Mechanism Underlying Successful Classification of Amnestic Mild Cognitive Impairment Using Multi-Sensory-Evoked Potentials

**DOI:** 10.1101/2024.08.10.607449

**Authors:** Lei Zhang, Malcom Binns, Ricky Chow, Rahel Rabi, Nicole D. Anderson, Jing Lu, Morris Freedman, Claude Alain

## Abstract

Early detection of amnestic mild cognitive impairment (aMCI) is crucial for timely interventions. This study combines scalp recordings of lateralized auditory, visual, and somatosensory stimuli with a flexible and interpretable support vector machine learning pipeline to differentiate individuals diagnosed with aMCI from healthy controls. Event-related potentials (ERPs) and functional connectivity (FC) matrices from each modality successfully predicted aMCI. Reduced ERP amplitude in aMCI contributed to classification. The analysis of FC using phase-locking value revealed higher FC in aMCI than controls in frontal regions, which predicted worse cognitive performance, and lower FC in posterior regions from delta to alpha frequency. We observe optimal classification accuracy (96.1%), sensitivity (97.7%) and specificity (94.3%) when combining information from all sensory conditions than when using information from a single modality. The results highlight the clinical potential of sensory-evoked potentials in detecting aMCI, with optimal classification using both amplitude and oscillatory-based FC measures from multiple modalities.

## Background

Amnestic mild cognitive impairment (aMCI) represents an intermediate stage between healthy aging and dementia, typically due to Alzheimer’s disease (AD), the most common cause of dementia in the aging population. Individuals with aMCI show impairment in episodic memory and sometimes in additional cognitive domains such as executive functioning and language,^1–3^ with preserved functional independence in daily activities.^4^ The diagnosis, prognosis, and management of aMCI remain challenging. Timely identification and intervention in aMCI are pivotal for its effective clinical management. The present study contributes to this critical need by introducing a cutting-edge framework that integrates the power of support vector machine (SVM) algorithms with a novel multisensory evoked potential (MSEP) paradigm.

Scalp-recording of event-related potentials (ERP) may help identify individuals with aMCI. Relative to healthy controls, individuals with MCI (including amnestic and non-amnestic subtypes that are prodromal to all-cause dementia) show significant reductions in N2 and P3 amplitude, as well as delayed P3 latency, signifying deficits in inhibitory control.^5–12^ Multivariate classification of cognitive ERPs has been shown to correctly discriminate participants with MCI from healthy controls with an accuracy ranging from 70% to 80% depending on the task.^13–15^ However, Cognitive ERP recordings can be time-consuming due to variability in amplitude and latency, requiring numerous trials for reliable signal-to-noise ratios. Additionally, cultural factors like language, education, and literacy can influence performance on cognitive ERP paradigms.^16^

To overcome these limitations, we developed the MSEP paradigm,^17^ which provides a means to delineate normal from impaired sensory processing. In this paradigm, lateralized auditory, visual, and somatosensory stimuli are presented sequentially at a high rate (e.g., 3-5 stimuli per second) and in random order. This approach minimizes neural adaptation, thereby ensuring reliable and fairly quick (∼15 minutes) measurements of sensory-evoked responses from three sensory modalities. Because these sensory-evoked responses are recorded while participants sit quietly in the absence of a task (i.e., no response required), there are no concerns related to task complexity, or difficulty understanding complex instructions, making it easy to administer to older adults. Therefore, the design of this paradigm provides practicality in clinical settings as this paradigm is short in length, accessible and inclusive for older adults, and can provide reliable measures of brain functioning.

The MSEP paradigm has been developed in light of prior ERP studies using either auditory,^5,18,19^ visual^20–22^ or somatosensory^23,24^ stimuli, which have revealed aberrant (hypersensitive and/or delayed responses) sensory-evoked responses in healthy older adults and those with aMCI. This over-activation of the sensory cortices may be linked to widespread structural changes in prefrontal regions, which may be exacerbated in individuals with MCI.^25–35^ Notably, lesions to prefrontal cortices have been associated with an increase in the amplitude of auditory, visual and somatosensory ERPs, which was attributed to deficits in inhibitory control.^36,37^ Moreover, sensory evoked response amplitude has been associated with deficits in memory tasks.^38–40^ Given these findings from studies measuring sensory evoked response amplitude from a single sensory modality, we foresee that using measurements from auditory, visual and somatosensory systems could significantly enhance the sensitivity of EEG measures in detecting aMCI.

Evidence suggests that SVM algorithms can successfully identify older adults with MCI (including amnestic and non-amnestic subtypes) from healthy controls with accuracy ranging from 75% to 98% using speech-evoked responses^41^ or resting-state EEG^42–44^ depending on the number and type of EEG features analyzed. A recent study applying parameter-optimized SVM classifiers to oscillatory-based functional connectivity data showed hearing status could be decoded during a speech categorization task solely using network-level descriptions.^45^ Extending prior work,^45^ we used adjusted connectome-based predictive modeling with SVM classifiers to distinguish older adults with a diagnosis of aMCI from healthy controls using EEG functional connectivity measures during the processing of auditory, visual, and somatosensory stimuli. Given prior findings of aberrant sensory-evoked responses in single sensory modalities, we hypothesize that the correct classification of aMCI will increase when combining information for multiple sensory modalities compared to using information from a single sensory modality.

## Methods

### Participants

Participants were recruited if they were native English speakers or learned English before the age of five, had a normal or corrected-to-normal vision (e.g., no history of degenerative conditions, glaucoma, cataracts significant enough to impede vision or color blindness), and reported no significant hearing loss, no history of learning disabilities, stroke, transient ischemic attack, traumatic brain injury with loss of consciousness greater than five minutes, substance abuse disorder, history of intracranial surgery, and any other diagnosis of major neurological or psychiatric disorder. Participants were excluded if they had a history of myocardial infarction, coronary artery disease, or bypass surgery. Participants were also excluded if they were prescribed hearing aids, taking medication known to possibly affect cognitive functioning, including antidepressants, anticonvulsants, antipsychotics, or consuming recreational drugs either concurrently or within the year before testing. An inclusion criterion for older adults was scoring above the cut-off on the Telephone Interview for Cognitive Status-Modified (TICS-M).^46^ Our final sample included 44 individuals with aMCI (62-88 years, 23 males) and 35 healthy older adult controls (60-88 years, 20 males). Participants were recruited from the Rotman Research Institute participant database. The Research Ethics Board of the Rotman Research Institute at Baycrest approved the study protocol. All participants provided informed written consent and were compensated a modest stipend for their participation and out-of-pocket expenses (e.g., parking).

### Neuropsychological assessment

Before recording neuroelectric activity, participants were administered a battery of standardized neuropsychological tests in the domains of intellectual functioning, memory, language, processing speed, and executive functioning. The neuropsychological assessment took place on a separate day from measurements of sensory-evoked responses to prevent fatigue effects. A registered neuropsychologist verified the diagnosis of single-domain or multiple-domain aMCIby. A full description of the administered neuropsychological assessments has been previously reported in Chow et al^47^ and Rabi et al.^48^ Demographic, neuropsychological, and clinical data for individuals with aMCI and age-matched controls are displayed in Table 1. The aMCI and control groups did not statistically differ in age, sex, or years of education.

**Table 1.**
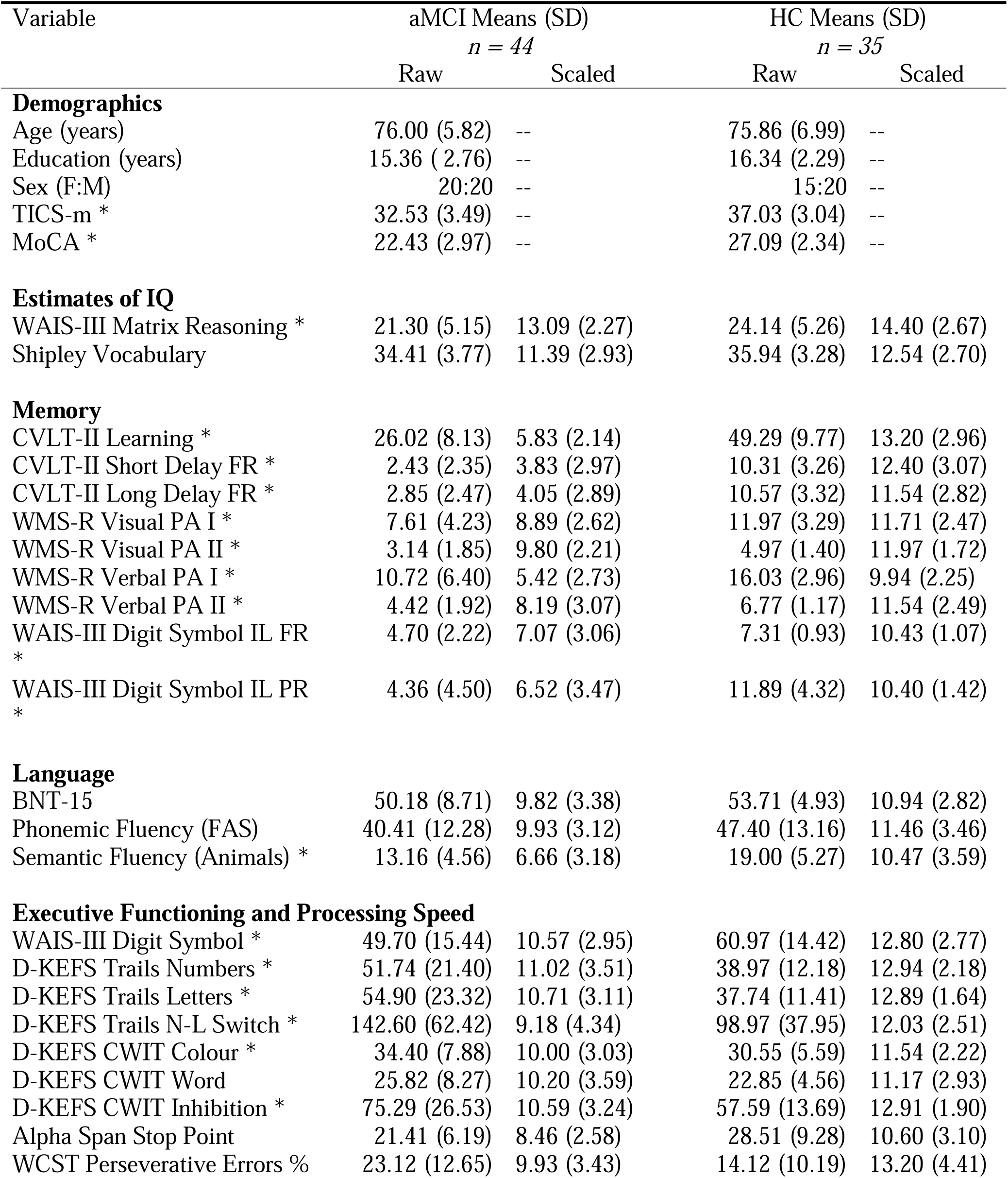
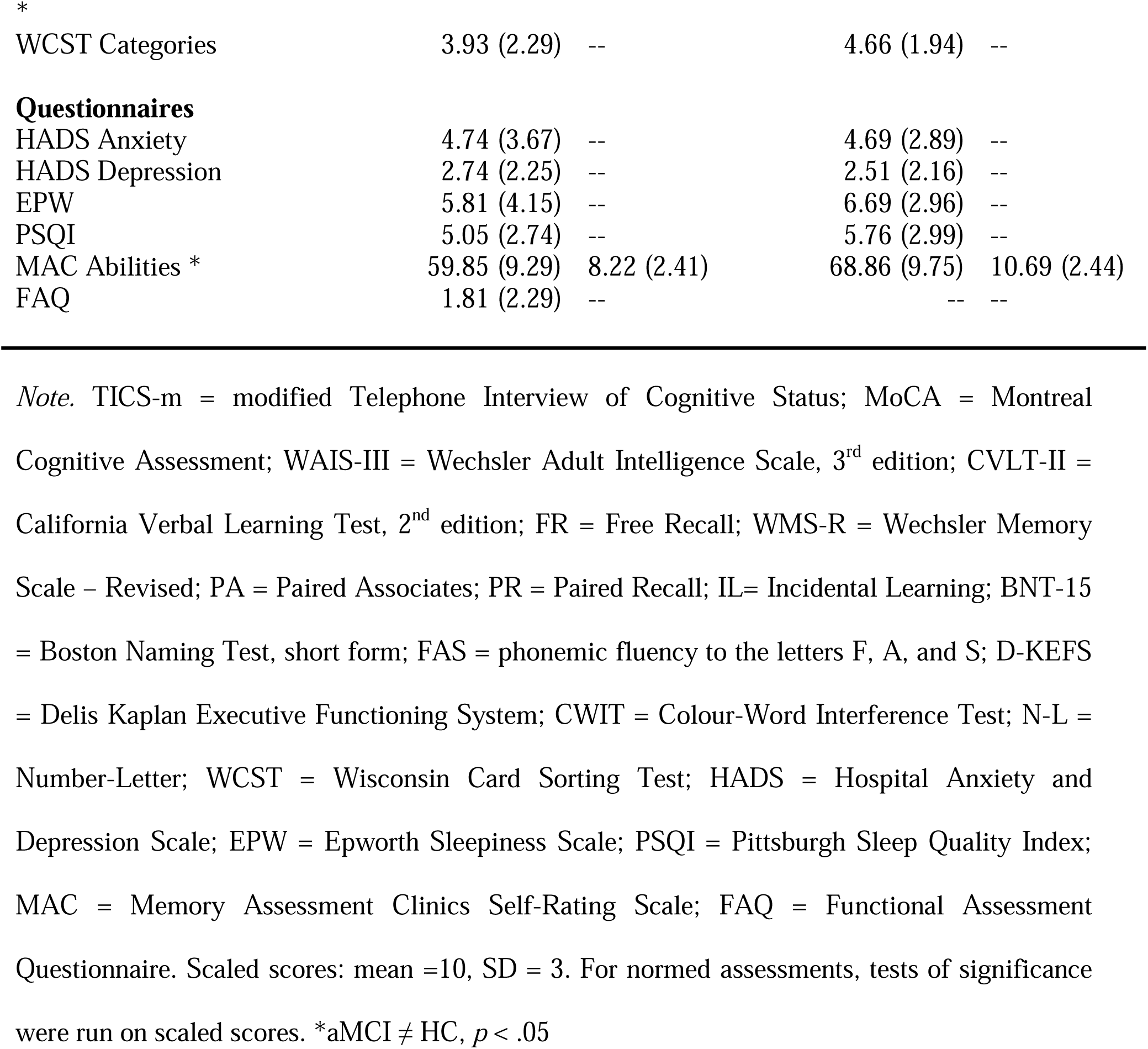
Participant Demographic and Neuropsychological Data.

| Variable | aMCI Means (SD)<br>*n* = 44 |  | HC Means (SD)<br>*n* = 35 |  |
| --- | --- | --- | --- | --- |
|  | Raw | Scaled | Raw | Scaled |
| **Demographics** |  |  |  |  |
| Age (years) | 76.00 (5.82) | -- | 75.86 (6.99) | -- |
| Education (years) | 15.36 ( 2.76) | -- | 16.34 (2.29) | -- |
| Sex (F:M) | 20:20 | -- | 15:20 | -- |
| TICS-m * | 32.53 (3.49) | -- | 37.03 (3.04) | -- |
| MoCA * | 22.43 (2.97) | -- | 27.09 (2.34) | -- |
| **Estimates of IQ** |  |  |  |  |
| WAIS-III Matrix Reasoning * | 21.30 (5.15) | 13.09 (2.27) | 24.14 (5.26) | 14.40 (2.67) |
| Shipley Vocabulary | 34.41 (3.77) | 11.39 (2.93) | 35.94 (3.28) | 12.54 (2.70) |
| **Memory** |  |  |  |  |
| CVLT-II Learning * | 26.02 (8.13) | 5.83 (2.14) | 49.29 (9.77) | 13.20 (2.96) |
| CVLT-II Short Delay FR * | 2.43 (2.35) | 3.83 (2.97) | 10.31 (3.26) | 12.40 (3.07) |
| CVLT-II Long Delay FR * | 2.85 (2.47) | 4.05 (2.89) | 10.57 (3.32) | 11.54 (2.82) |
| WMS-R Visual PA I * | 7.61 (4.23) | 8.89 (2.62) | 11.97 (3.29) | 11.71 (2.47) |
| WMS-R Visual PA II * | 3.14 (1.85) | 9.80 (2.21) | 4.97 (1.40) | 11.97 (1.72) |
| WMS-R Verbal PA I * | 10.72 (6.40) | 5.42 (2.73) | 16.03 (2.96) | 9.94 (2.25) |
| WMS-R Verbal PA II * | 4.42 (1.92) | 8.19 (3.07) | 6.77 (1.17) | 11.54 (2.49) |
| WAIS-III Digit Symbol IL FR * | 4.70 (2.22) | 7.07 (3.06) | 7.31 (0.93) | 10.43 (1.07) |
| WAIS-III Digit Symbol IL PR * | 4.36 (4.50) | 6.52 (3.47) | 11.89 (4.32) | 10.40 (1.42) |
| **Language** |  |  |  |  |
| BNT-15 | 50.18 (8.71) | 9.82 (3.38) | 53.71 (4.93) | 10.94 (2.82) |
| Phonemic Fluency (FAS) | 40.41 (12.28) | 9.93 (3.12) | 47.40 (13.16) | 11.46 (3.46) |
| Semantic Fluency (Animals) * | 13.16 (4.56) | 6.66 (3.18) | 19.00 (5.27) | 10.47 (3.59) |
| **Executive Functioning and Processing Speed** |  |  |  |  |
| WAIS-III Digit Symbol * | 49.70 (15.44) | 10.57 (2.95) | 60.97 (14.42) | 12.80 (2.77) |
| D-KEFS Trails Numbers * | 51.74 (21.40) | 11.02 (3.51) | 38.97 (12.18) | 12.94 (2.18) |
| D-KEFS Trails Letters * | 54.90 (23.32) | 10.71 (3.11) | 37.74 (11.41) | 12.89 (1.64) |
| D-KEFS Trails N-L Switch * | 142.60 (62.42) | 9.18 (4.34) | 98.97 (37.95) | 12.03 (2.51) |
| D-KEFS CWIT Colour * | 34.40 (7.88) | 10.00 (3.03) | 30.55 (5.59) | 11.54 (2.22) |
| D-KEFS CWIT Word | 25.82 (8.27) | 10.20 (3.59) | 22.85 (4.56) | 11.17 (2.93) |
| D-KEFS CWIT Inhibition * | 75.29 (26.53) | 10.59 (3.24) | 57.59 (13.69) | 12.91 (1.90) |
| Alpha Span Stop Point | 21.41 (6.19) | 8.46 (2.58) | 28.51 (9.28) | 10.60 (3.10) |
| WCST Perseverative Errors % | 23.12 (12.65) | 9.93 (3.43) | 14.12 (10.19) | 13.20 (4.41) |

\*

|  |  |  |  |  |
| --- | --- | --- | --- | --- |
| WCST Categories | 3.93 (2.29) | -- | 4.66 (1.94) | -- |
| **Questionnaires** |  |  |  |  |
| HADS Anxiety | 4.74 (3.67) | -- | 4.69 (2.89) | -- |
| HADS Depression | 2.74 (2.25) | -- | 2.51 (2.16) | -- |
| EPW | 5.81 (4.15) | -- | 6.69 (2.96) | -- |
| PSQI | 5.05 (2.74) | -- | 5.76 (2.99) | -- |
| MAC Abilities * | 59.85 (9.29) | 8.22 (2.41) | 68.86 (9.75) | 10.69 (2.44) |
| FAQ | 1.81 (2.29) | -- | -- | -- |

*Note.* TICS-m = modified Telephone Interview of Cognitive Status; MoCA = Montreal Cognitive Assessment; WAIS-III = Wechsler Adult Intelligence Scale, 3<sup>rd</sup> edition; CVLT-II = California Verbal Learning Test, 2<sup>nd</sup> edition; FR = Free Recall; WMS-R = Wechsler Memory Scale – Revised; PA = Paired Associates; PR = Paired Recall; IL= Incidental Learning; BNT-15 = Boston Naming Test, short form; FAS = phonemic fluency to the letters F, A, and S; D-KEFS = Delis Kaplan Executive Functioning System; CWIT = Colour-Word Interference Test; N-L = Number-Letter; WCST = Wisconsin Card Sorting Test; HADS = Hospital Anxiety and Depression Scale; EPW = Epworth Sleepiness Scale; PSQI = Pittsburgh Sleep Quality Index; MAC = Memory Assessment Clinics Self-Rating Scale; FAQ = Functional Assessment Questionnaire. Scaled scores: mean =10, SD = 3. For normed assessments, tests of significance were run on scaled scores. \*aMCI ≠ HC, $p < .05$

### Stimuli and multi-sensory evoked paradigm

The visual stimulus was a 5 by 10 square black and white checkerboard pattern presented at 50ms in duration that spanned the entire height and approximately 40% of the width of a 19-inch computer monitor that extended from either the left or right edge of the monitor. The entire checkerboard stimulus subtended a visual angle of 14.3° by 28.1°, with each checkerboard square subtending a visual angle of 2.9° by 2.9°. The auditory stimulus was a monoaural harmonic complex tone (50 ms in duration, 5 ms rise/fall time). It comprised five tonal elements (200, 400, 600, 800, and 1000 Hz) and had a fundamental frequency of 200 Hz. The tone was presented to the left or right ear through ER-3A insert earphones (Etymotic Research Inc., Elk Grove, IL, USA). The sound intensity was set at 85 decibels (dB) sound pressure level (SPL) as measured by a Larson Davis 824 SPL meter using a 2cc coupler. The somatosensory stimulus consisted of pneumatic tactile stimulation delivered simultaneously to the tips of all four fingers (thumb excluded) of the left or right hand. The somatosensory stimulation was delivered from a puff-type pressure stimulator from a compressed air tank with electromagnetic air valves kept constant at 30 psi through plastic tubes to 8 pneumatically driven inflatable circular plastic membranes with a diameter of 1 cm each. Each membrane was worn on the participant’s fingers at the distal interphalangeal joint. During somatosensory stimulation, the valves were activated to periodically inflate the membrane and deliver brief pressure pulses of 50 ms duration to the fingertips.

### Procedure

Participants were presented with lateralized visual, auditory, and somatosensory stimuli sequentially in a randomized order while sitting in a sound-attenuated booth. Before recording sensory evoked potentials, participants were presented with a 10-second sample of the paradigm to ensure they could clearly perceive the stimuli. During the stimulus presentation, participants were instructed to gaze on a fixation dot 0.60 cm in diameter at the center of a 19-inch computer screen set at 100% contrast level and placed 60 cm from participants. The stimulus onset asynchrony varied between 200 and 340 ms (20 ms steps, rectangular distribution). The stimulus presentation lasted approximately 12 minutes, with 1440 stimuli (480 trials per sensory modality) evenly distributed between left and right laterality. Stimuli were presented using Presentation software (version 13, Neurobehavioral Systems, Albany, CA).

### EEG Acquisition and Preprocessing

Neuroelectric brain activity was recorded continuously using a 76-channel BioSemi ActiveTwo acquisition system (BioSemi, Amsterdam, The Netherlands) with a sampling rate of 512 Hz. Sixty-six electrodes were positioned on the scalp using a BioSemi headcap according to the standard 10-20 system with a pair of Common Mode Sense active electrode and Driven Right Leg passive electrode serving as ground. Ten additional electrodes were placed below the hairline (i.e., mastoids, pre-auricular points, two lateral ocular sites, two inferior ocular sites, and two additional frontal-lateral electrodes) to monitor eye movements and cover the whole scalp evenly. EEG recordings were preprocessed offline using FieldTrip toolbox^49^ and custom scripts.

The EEG data were first re-referenced to the average of all electrodes. Then, independent component analysis was performed to remove the artifacts caused by eye blinks, eye movements, and heartbeat. Next, the EEG data were bandpass filtered from 1 to 40Hz. Filtered data were parsed into 500 ms epochs time-locked to stimulus onset, including 100 ms of pre-stimulus activity and baseline corrected using pre-stimulus interval (-100 to -20 ms). Lastly, the preprocessed data were visually inspected to identify trials contaminated with artifacts and noisy electrodes; these trials were then rejected from subsequent analysis. Noisy electrodes were interpolated, with no more than 10% of the channels interpolated per participant.

We did not have a priori hypotheses regarding possible stimulus lateralization. Hence, as a dimension reduction method and to improve the signal-to-noise ratio, epochs of lateralized sensory-evoked potentials were averaged together in a transposed montage such that right-lateralized responses (elicited by left-lateralized stimuli) were kept untransposed, and left-lateralized evoked potentials (elicited by right-lateralized stimuli) were transposed across laterality. This transposition method allowed us to combine the evoked responses from lateralized stimuli, significantly increasing the number of trials per modality.

### Electrode-level functional connectivity matrix construction

A surface Laplacian method was applied to reduce the effect of volume conduction on functional connectivity estimates.^50^ The surface Laplacian is a spatial filter that computes the second spatial derivative of the scalp potential, which provides a measure of the local current source density. Then, for each sensory modality, phase locking value (PLV), which estimated the phase synchronization of two signals, was calculated for estimating the functional connectivity between pairs of electrodes across 12 frequency bands ranging from 2.5 Hz to 30 Hz, with each band being 2.5 Hz in width (i.e., 2.5 Hz, 5 Hz, 7.5 Hz … 27.5 Hz, 30 Hz). Consequently, for each participant, twelve 72 by 72 functional connectivity matrices were created for each stimulus modality per participant.

### Adjusted connectome-based predictive modeling (CPM)

We adjusted the procedure from CPM^51^ and used it with SVM learning to classify each participant as belonging to either the aMCI or control group (Fig. 1). Functional connectivity matrices of each modality and frequency band were separately entered into the classification analysis. Leave-one-subject-out (LOSO) cross-validation was employed to evaluate classification performance (i.e., classification accuracy) for each electrode under each sensory condition. Specifically, for each cross-validation fold, matrices from 78 participants were used to build the model and the matrix from the left-out participant was used to test model performance. For each matrix, we first performed a two-sample independent *t*-test (two-tailed) on each edge between aMCI and healthy control groups using the training set. Edges that showed sufficient group difference (p_uncorrected_ < 0.05) were selected for training the SVM classification model with a linear kernel. A parameter search was performed using training data to optimize the cost value, which balanced the margin and training error for each model and each stimulus modality (thereby maximizing the average accuracy of the top three frequency bands over the range 2^-6^, 2^-5^,…, 2^5^). After training the model, the edges from the left-out participant were used to test the model. In the model training process, the left-out data is not involved in the model training and hyperparameter selection to avoid over-fitting problems. With the LOSO cross validation procedure, every participant would be predicted to be in the aMCI or control group using the model trained by the selected functional connectivity edges of all other participants. Classification accuracy was calculated by averaging the accuracy across all folds. SVM classification analysis was conducted using the LIBSVM package.^52^ The selected edge masks in each fold were kept for visualizing the important edges that contributed to the classification.

**Fig. 1.**
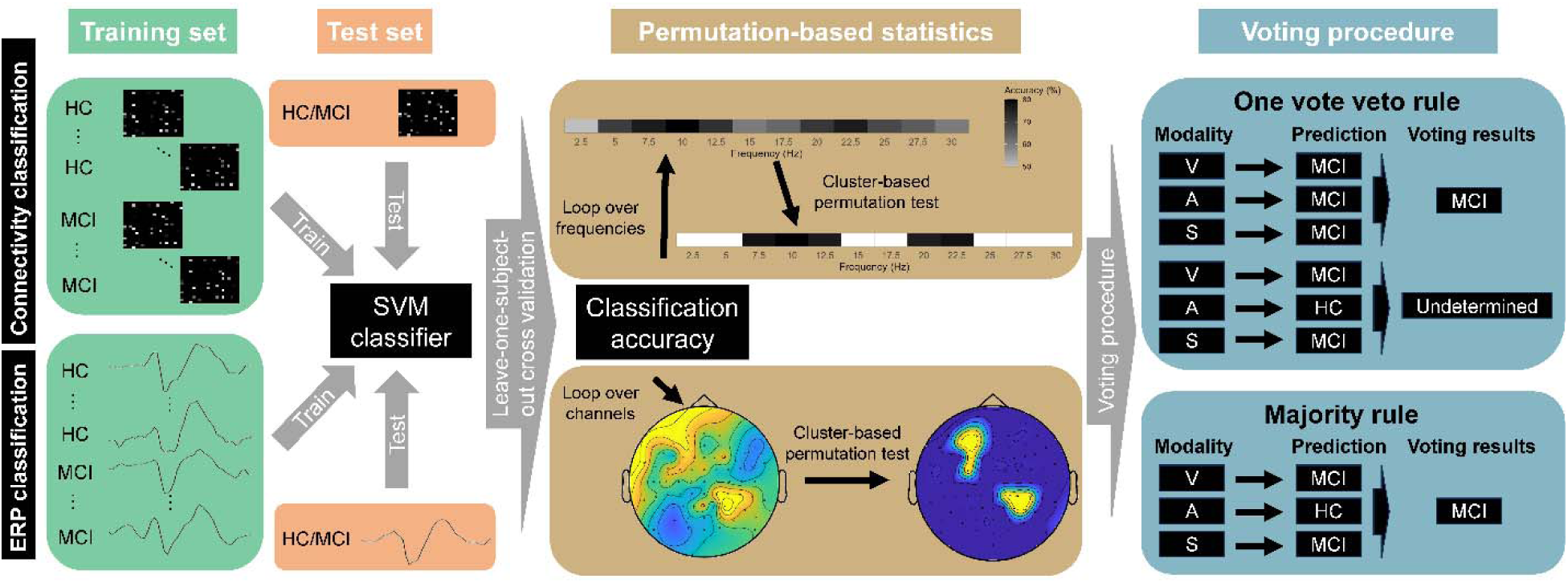
Classification procedure illustration. We separately performed SVM classification analyses using functional connectivity matrices or ERP for each sensory modality (for details, please see methods). With leave-one-subject-out cross validation, we obtain classification accuracy of each frequency or electrode for each sensory modality. Next, we performed permutation-based test for the significance and statistics of each frequency or electrode for each sensory modality (for details, please see methods). Lastly, we employed the voting procedures to combine the output from difference neural indices evoked by stimuli from difference sensory modalities to perform the prediction.

We performed permutation testing to determine the statistical significance of the classification accuracy of each frequency band for each sensory modality. The participants were shuffled to perform the classification analysis described above. The number of permutations was set at 2000, allowing us to construct the null distributions of classification accuracy for each frequency band under three sensory modalities. Permutation-based statistics were calculated based on the following formula:

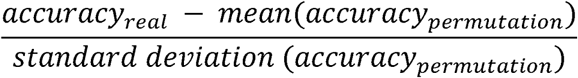

The frequency-level permutation-based statistics were followed by cluster-based permutation tests to correct for multiple comparisons across frequency bands. The sums of the permutation-based statistics of contiguous frequencies that were significant (*p_permutation_* < 0.05) were computed. Then, 2000 permutations were conducted, and the largest permutation-based statistics mass for each permutation was extracted to get the null distribution of the cluster-level mass. Significance was set at a family-wise corrected *p*-value of *p_fwe_* < 0.05 cluster-mass corrected.

### SVM classification with sensory-evoked response amplitude

Besides functional connectivity matrices, we also performed SVM classification analysis using the amplitude of sensory-evoked potentials (Fig. 1). Averaged amplitudes for each modality were separately entered in the classification analysis. For each electrode, classifiers were trained using the SVM algorithm with a linear kernel separately for the three modalities. Similarly, LOSO cross-validation was employed to evaluate classification performance (i.e., classification accuracy) for each electrode in each sensory modality. A parameter search was also performed using training data to optimize the cost value for each model and each stimulus modality (thereby maximizing the average accuracy of the top ten electrodes over the range 2^-3^, 2^-2^,…, 2^5^). We used cluster-based permutation testing to determine the statistical significance of the classification accuracy of each electrode for each modality. Channel neighbors were defined by the FieldTrip function “ft_prepare_neighbours” with the “triangulation” method.^49^ We computed the sums of the permutation-based statistics of contiguous channels that were significant (*p_permutation_* < 0.05). Then, 2000 permutations were conducted, and the largest permutation-based statistics mass for each permutation was extracted to get the null distribution of the cluster-level mass. Significance was set at *p_fwe_* < 0.05 cluster-mass corrected.

### Voting method for classification combining multiple modalities

We used voting strategies to combine the classification results from different modalities and indices. The frequency bands or electrodes with the highest classification accuracy in each sensory modality were selected and entered into the voting procedure. We employed two strategies in the voting procedure: majority rule or one-vote veto rule. The majority rule denotes that whether a given participant is classified as belonging to the aMCI or control group is determined by most supporters (i.e., a participant was classified as belonging to one group if they were also identified as belonging to the same group in two or all three modalities). As for the one-vote veto rule, a given participant was classified as belonging to the aMCI or control group only when all involved classifiers agreed on aMCI status. The left-out participants were labelled as undetermined and not included in the accuracy calculation.

### Group comparison of evoked response from three modalities

Averaged sensory-evoked responses in the significant clusters under each modality were entered into the group comparison analysis between aMCI and control groups. Note that the participants with more than one channel that misclassified them were not included in the group comparison since they could not be classified accurately across channels. Independent-sample *t-* tests compared the group-averaged sensory-evoked response amplitudes among significant clusters between aMCI and controls. Cluster-based permutation testing was performed to find clusters that showed significant group differences between the two groups.

### Brain-behavior correlation

Linear regression models were fit to examine the relationship between neural indices (functional connectivity estimates of the important edges, sensory-evoked response amplitude of significant clusters) and composite scores from the neuropsychological assessment (Table 1). Group variable (aMCI/healthy older adults) was set as the control variable in the linear regression models.

## Results

### Classification with unisensory prediction models

With the adjusted CPM approach, we trained the SVM learning classifiers to predict whether a given participant belonged to the aMCI or control group using unisensory functional connectivity matrices (visual, auditory, or somatosensory). We found frequency clusters that showed significantly greater classification accuracy than chance for all three sensory modalities (Fig. 2A). For visual stimuli, functional connectivity matrices from 2.5 Hz to 10 Hz and from 15 Hz to 17.5 Hz significantly predicted correct classification (i.e., whether a participant belonged to the aMCI or healthy control group, 2.5Hz-10Hz: *p_fwe_* = 0.007, 15 Hz-17.5 Hz: *p_fwe_* = 0.040). For auditory stimuli, the significant frequency bands were from 2.5 Hz to 7.5 Hz and 12.5 Hz (2.5 Hz-7.5 Hz: *p_fwe_* < 0.001, 12.5 Hz: *p_fwe_* < 0.001). For somatosensory stimuli, the significant frequency bands were from 2.5 Hz to 12.5 Hz (*p_fwe_* < 0.001).

**Fig. 2.**
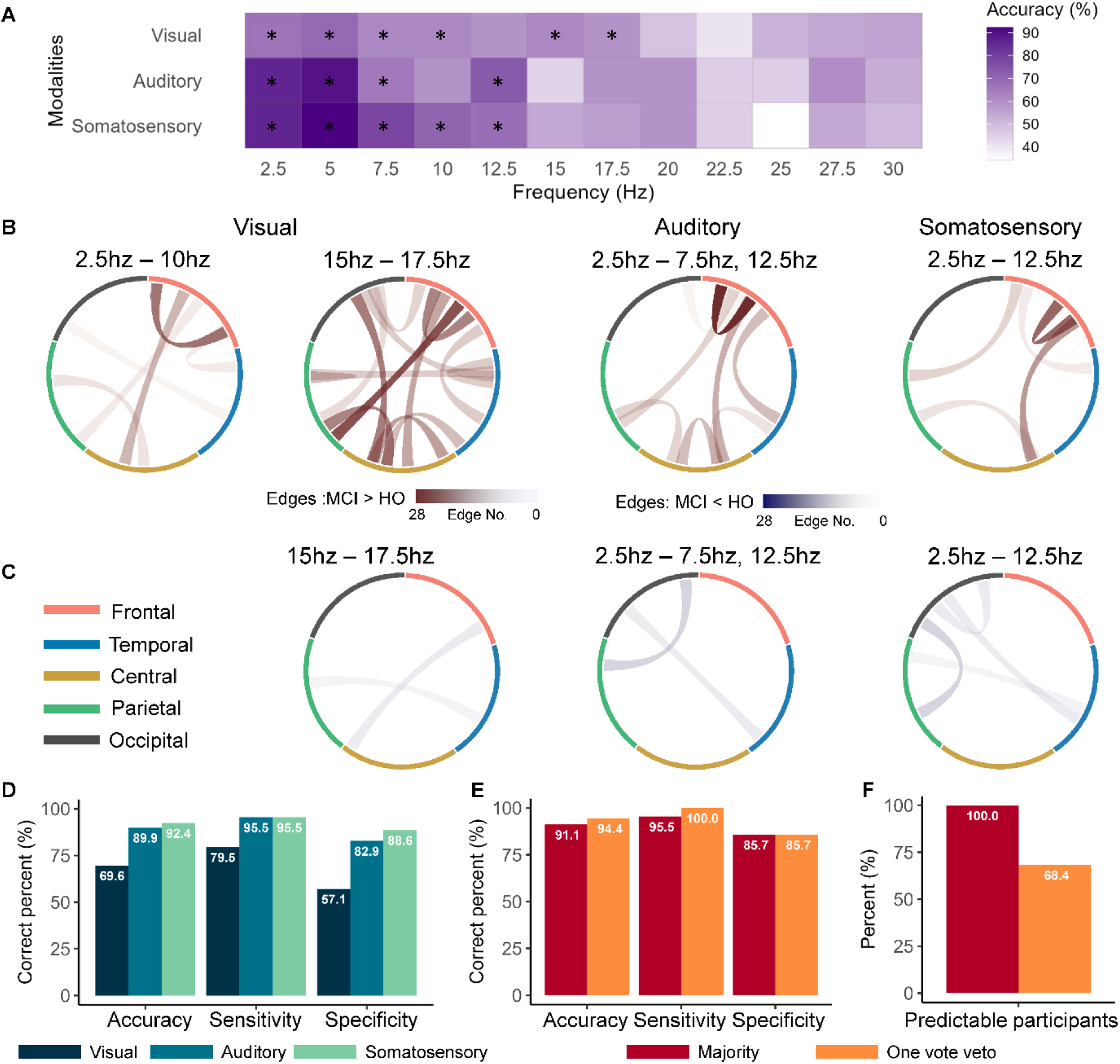
Classification results using functional connectivity matrices. A. Classification accuracy at different frequency bands for three modalities. *denotes the significant frequencies for each modality after cluster-based correction. B-C, Summarized the positive (B) and negative (C) important edge numbers that were kept according to the five ROIs (frontal ROI: frontopolar, anterior-frontal and frontal channels, temporal ROI: frontotemporal and temporal-parietal channels, central ROI: frontocentral and central channels, parietal ROI: centroparietal and parietal channels, and occipital ROI: parietal-occipital and occipital channels), Important connectivity edges were those edges that preserved in more than 90% folds. D-F, Classification accuracy, sensitivity, and specificity of the top frequencies (D), voting results combining three modalities (E), predictable participant percent (F).

Figure 2A shows classification accuracy per frequency band. Classification accuracies tended to be higher for lower frequencies (i.e., delta, theta, and alpha) in all three modalities, with the highest classification accuracy at 5 Hz (Fig. 2D, 69.6% for visual stimuli, *p_permutation_* = 0.004, 89.9% for auditory stimuli, *p_permutation_* < 0.001, 92.4% for somatosensory stimuli, *p_permutation_* < 0.001). The sensitivities of all three modalities were high, while the specificities were relatively low (Fig. 2D, for visual stimuli, 79.5% vs. 57.1%; for auditory stimuli, 95.5% vs. 82.9%; for somatosensory stimuli, 95.5% vs. 88.6%). In other words, the fitted model showed good detection of participants with aMCI (i.e., better able to detect true positives) but poor detection of healthy older adults (i.e., resulting in a few false positives).

We employed a voting method to combine the predictive power from three modalities. With the majority rule, the accuracy, sensitivity, and specificity of the model were similar to the somatosensory modality, which showed the best performance (i.e., Fig. 2E, Accuracy: 91.1%; Sensitivity: 95.5%; Specificity: 85.3%). The classification accuracy was calculated based on the subset of participants for whom predictions from all involved classifiers were consistent. There was an accuracy-proportion of predictable participant trade-off with one-vote veto rule. We observed a slight performance increase compared to the somatosensory classifier, but only 68.4% of participants could be accurately identified (Fig. 2E, F, Accuracy: 94.4%; Sensitivity: 100%; Specificity: 85.7%). The specificity was still relatively low with a voting procedure that combined the three modalities.

### Functional connectivity strength predicted cognitive performance

We extracted all of the masks used for each fold. Since LOSO cross-validation was used in SVM classification, 79 masks were generated in the analysis. Masks were slightly different for each fold because training sets varied for each fold. We kept the edges that were preserved in most folds (> 90%), which were more important for the classification analysis than the rest of the edges.

We divided the electrode montage into five regions of interest (ROIs): frontal (frontopolar, anterior-frontal and frontal channels), temporal (frontotemporal and temporal-parietal channels), central (frontocentral and central channels), parietal (centroparietal and parietal channels), and occipital (parietal-occipital and occipital channels). Then, we summarized the positive and negative edge numbers that were kept according to the five ROIs separately for each sensory modality and significant frequency band. Note that positive edges refer to the edges where aMCI showed higher functional connectivity estimates than healthy older adults. In comparison, negative edges refer to the edges where aMCI showed lower functional connectivity estimates.

Within lower frequency bands (i.e., delta, theta, and alpha bands), we found similar connectivity pattern for the three modalities (Fig. 2B and 2C). That is, the aMCI group showed stronger functional connectivity than healthy older adults within frontal electrodes and connectivity from frontal and central electrodes to posterior electrodes. Meanwhile, within the beta band, we also found that aMCI showed greater functional connectivity than controls across the whole brain with visual stimuli. Besides the increased connectivity, we also observed weaker edges in the aMCI group that connected posterior regions like occipital, temporal, and parietal areas within lower frequency bands. We found increased frontal-central and temporal-parietal connectivity within beta band in the aMCI group.

Linear regression analysis was performed to investigate the relationship between functional connectivity of the important edges and cognitive performance from neuropsychological assessments. We separately summed the important positive or negative edges for three modalities as the independent variable in the linear regression analysis. Group variable was inputted to the model as the control variable. We found that functional connectivity that was higher in the aMCI group than in healthy older adults predicted worse cognitive performance within both frequency bands (Table 2, EF: Delta-Theta-alpha band: Visual: β = - 4.37, p = 0.005; Somatosensory: β = -3.60, p = 0.017; Beta band: Visual: β = -5.39, p = 0.010). The higher functional connectivity in aMCI may reflect a loss of complexity in EEG dynamics, which has been associated with difficulties in adapting to new situations and could be a potential biomarker for aMCI diagnosis.^53–55^ However, for functional connectivity that was lower in the aMCI group, we observed different results. Within the beta band, weaker functional connectivity between frontal and posterior areas predicted better performance on language tests (Table 2, Language: β = -5.92, p = 0.006).

**Table 2.** Linear regression results. Regression coefficients and corresponding p values (in parenthesis) between summed functional connectivity and cognitive performance after controlling group.

|  | Edge Strength | Memory | Speed | EF | Language |
| --- | --- | --- | --- | --- | --- |
| Delta-Theta-Alpha band | V: aMCI > HC | 0.03<br>(0.982) | -2.09 (0.099) | -4.37 (0.005) | -2.07 (0.264) |
|  | A: aMCI > HC | 1.00 (0.554) | -1.57 (0.306) | -3.57 (0.063) | -2.38 (0.284) |
|  | S: aMCI > HC | -0.28 (0.833) | -1.18 (0.328) | -3.60 (0.017) | -1.74 (0.321) |
|  | A: aMCI < HC | 2.13 (0.243) | 0.50 (0.764) | 1.19 (0.569) | 0.34 (0.889) |
|  | S: aMCI < HC | 1.88 (0.272) | 2.44 (0.115) | 2.68 (0.172) | 1.29 (0.571) |
| Beta band | V: aMCI > HC | -2.71 (0.143) | -3.17 (0.058) | -5.39 (0.010) | -3.65 (0.135) |
|  | V: aMCI < HC | -0.01 (0.993) | -2.15 (0.154) | -2.55 (0.180) | -5.92 (0.006) |

*Note:* EF = executive functioning; HC = healthy control;

### Classification with unisensory evoked response amplitude

We also trained the SVM classifiers to predict whether the participant belonged to the aMCI or control group using the amplitude of sensory-evoked responses (visual, auditory, or somatosensory stimuli). Compared to a null distribution of classification accuracy generated by permutation testing, we found significant clusters that showed greater classification accuracy than null distribution for all three sensory modalities (Fig. 3A-C). For visual stimuli, we observed clusters that showed significant classification accuracy in left central-frontal (FP1, AF7, AF3, F3, FC3, *p_fwe_* = 0.003) and right parietal areas (CP6, CP4, P6, CP2, P4, *p_fwe_* = 0.001). For auditory stimuli, we found significant clusters in bilateral central areas (FC5, FC3, C1, Cz, C2, *p_fwe_* < 0.001), left temporal-parietal areas (P7, PO7, PO9, TP9, *p_fwe_* = 0.020, and right parietal areas (P6, P8, PO8, PO4, O2, *p_fwe_* < 0.001). For somatosensory stimuli, there was a significant cluster in the right occipital region (Iz, O2, *p_fwe_* = 0.040).

**Fig. 3.**
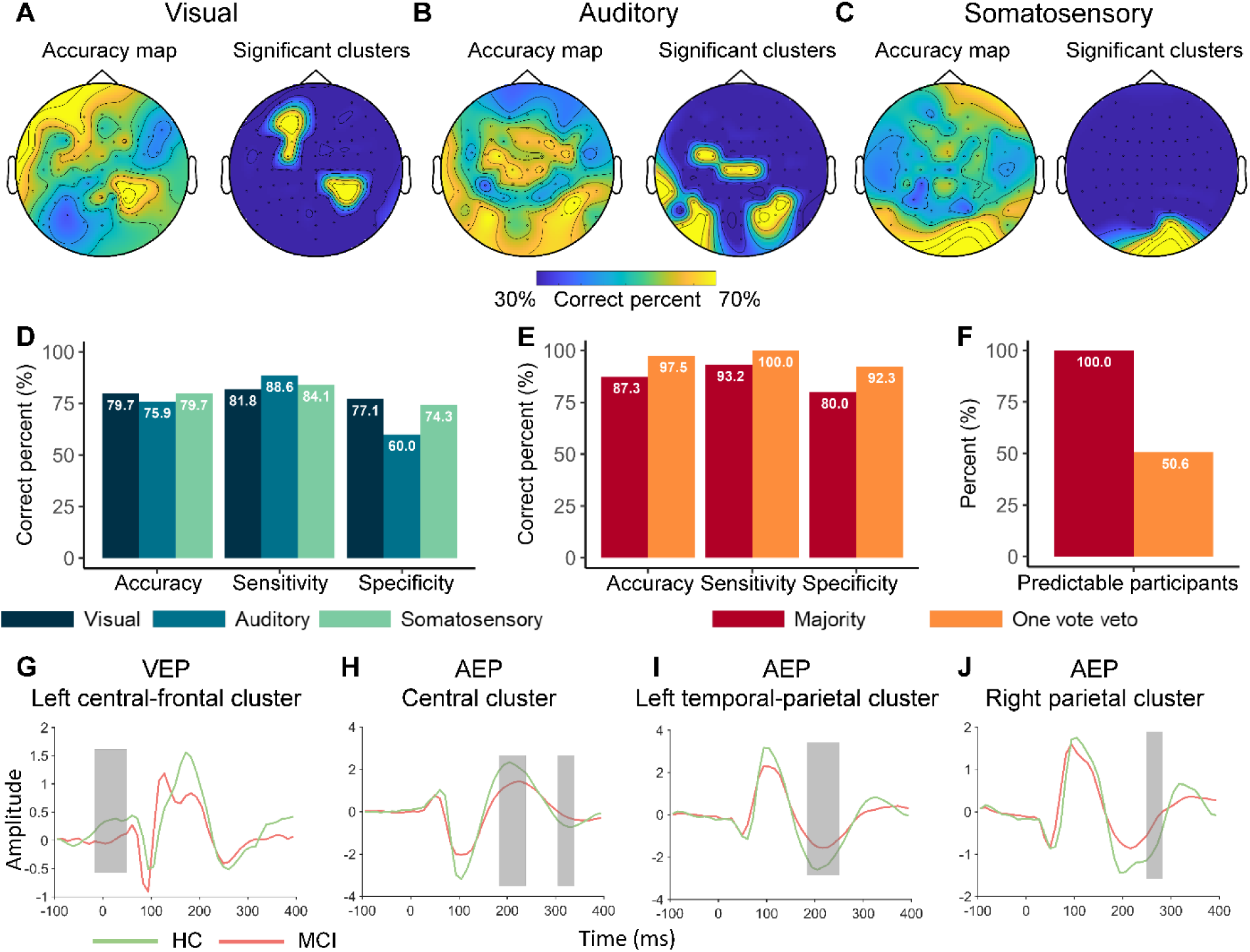
Classification results using ERP. A-C, Topoplot (left: unthresholded accuracy map; right: significant clusters) of prediction accuracy for visual (A), auditory (B), and somatosensory (C) modalities. D-F, Classification accuracy, sensitivity, and specificity of the top electrodes (D), voting results combining three modalities (E), predictable participant percent (F). G-J, Averaged ERP of the significant clusters.

Fig. 2D shows the electrode with the highest classification accuracy for visual (P8, classification accuracy = 79.7%; sensitivity = 81.8%; specificity = 77.1%), auditory (P4, classification accuracy = 76.0%; sensitivity = 88.6%; specificity = 60.0%), and somatosensory stimuli (Iz, classification accuracy = 79.7%; sensitivity = 84.1%; specificity = 74.3%). Again, similar to the model trained by functional connectivity matrices, the model’s specificity is lower than the sensitivity.

Models trained with unisensory evoked response amplitude were not as good as models trained with unisensory functional connectivity. However, as for EEG connectivity, model performance increased significantly after combining three modalities using the voting procedure. With majority rule among classifiers of all three modalities, classification accuracy increased to 87.3% (Fig. 3E, accuracy = 87.3%, sensitivity = 93.2%, specificity = 80.0%). With the one-vote veto rule, the model performance slightly increased compared to the model trained by functional connectivity. Classification accuracy increased to 97.5% (Fig. 3E-F, sensitivity = 100%, specificity = 92.3%) at the expense that only 50.6% of participants could be predicted.

### Reduced sensory evoked response amplitude in aMCI

To unravel what contributes to the predictability of the SVM classifier trained by averaged evoked responses from the three sensory modalities, we compared the averaged sensory-evoked response amplitude among the significant clusters between the aMCI group and healthy older adults. Only clusters that contained no more than one electrode that generated false predictions were included.

For visual stimuli, we found a larger early positive component over the frontocentral region ipsilateral to stimulus presentation in healthy older adults than those with aMCI (Fig. 3G,from -28ms to 50ms, *p_fwe_* = 0.003). However, we did not find a significant difference in contralateral parietal areas. For auditory stimuli, older adults showed greater positive-negative evoked response amplitude at about 200 and 300 ms after sound onset, referred to as P2 and N3, respectively (Fig. 3H, P2: from 171ms to 238ms, *p_fwe_* = 0.004; N3: from 294ms to 338ms, *p_fwe_* = 0.004) in central cluster, greater P2 amplitude (Fig. 3I, from 172ms to 249ms, *p_fwe_* = 0.002) in a left temporal-parietal cluster, and greater P2 amplitude (Fig. 3J, from 238ms to 283ms, *p_fwe_* = 0.016) in right parietal cluster. For somatosensory stimuli, there was no significant difference between the two groups.

Next, we used a linear regression to investigate the relationship between significant clusters and cognitive performance in older adults. After controlling for the group as the control variable, we found that a greater auditory evoked N3 amplitude in central regions significantly predicted higher speed composite scores (*β_AEP_* = -0.36, p = 0.029).

### Combining classification models trained by functional connectivity and ERPs

We combined the six models (3 modalities × 2 neural indices) with voting procedure because both models trained with connectivity measures and ERP amplitude (after combining three modalities) were relatively low with the majority rule. With the one-vote veto rule, only 35.4% of participants could be predicted; therefore the model showed little utility. With the majority rule, classification accuracy was 96.1%, and we achieved balanced sensitivity and specificity (Fig. 4A-B, sensitivity = 97.7%, specificity = 94.3%).

**Fig. 4.**
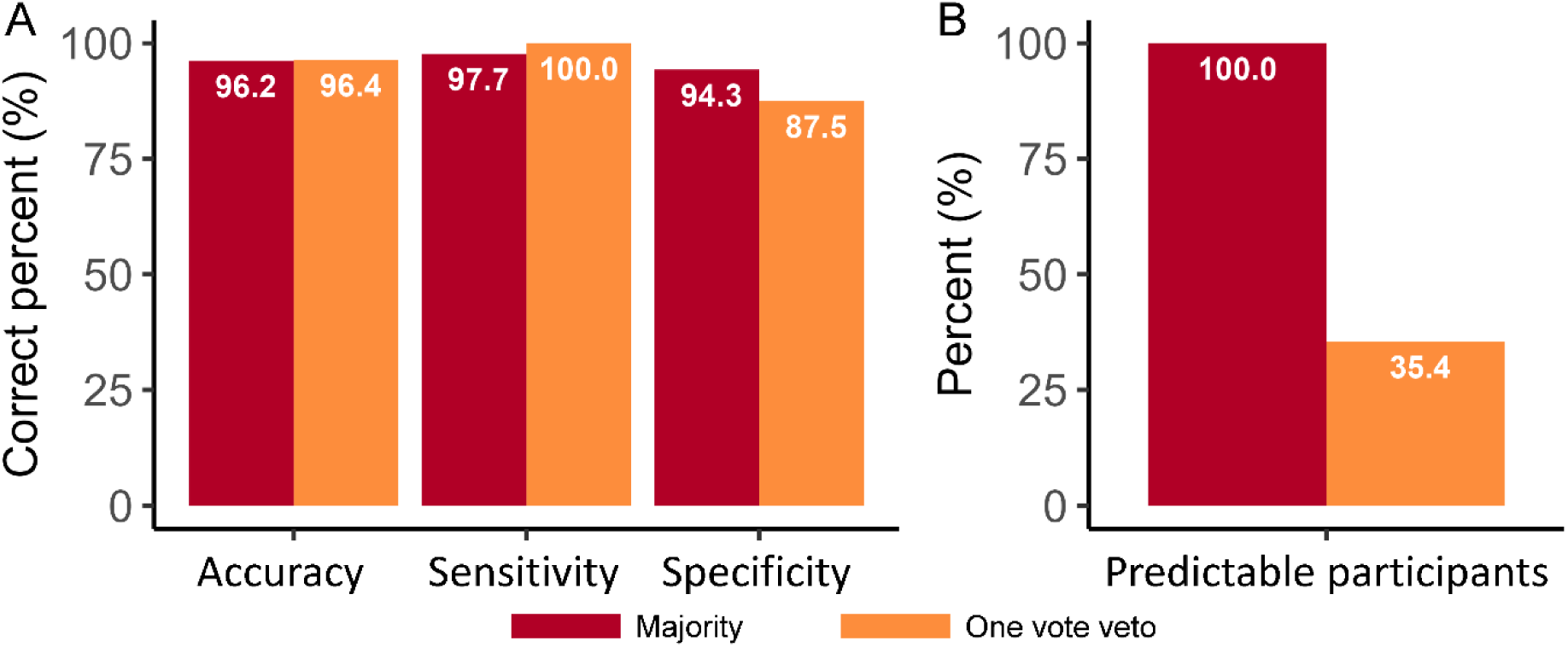
Voting results combining six models. We employed majority rules and one vote veto rules to make predictions based on the predictions from the models trained by functional matrices or ERP evoked by visual, auditory, and somatosensory modalities. A-B, Classification accuracy, sensitivity, and specificity of voting results combining three modalities (A), predictable participants percent (B).

## Discussion

This study employed the SVM algorithm to classify individuals with aMCI and healthy older adults with electrode-wise functional connectivity matrices and sensory-evoked responses from auditory, visual, and somatosensory systems. We found that healthy older adults and those with aMCI could be reliably classified using neural indices for each sensory modality. However, we observed a significant increase in classification accuracy after combining both neural indices under all three sensory conditions with the majority rule voting procedure, which yielded optimal classification accuracy (96.1%), sensitivity (97.7%), and specificity (94.3%). With adjusted CPM analysis, we identified increased functional connectivity within frontal and central regions in a lower frequency band and whole-brain functional connectivity in the beta band, the major contributing features in classification. Meanwhile, a group comparison of sensory-evoked response amplitude between correctly classified participants with aMCI and healthy older adults showed smaller response amplitude in the aMCI group than healthy older adults at central, frontal, parietal, and temporal electrodes. Notably, these increases and reductions in connectivity were correlated with better cognitive performance, namely executive functioning and speed, suggesting these features could be sensitive to cognitive impairment often associated with aMCI.

Since individuals with aMCI often experience difficulty with complex decision-making, experimental paradigms that are easy, fast, and require low cognitive demands are needed for clinical assessment of aMCI. Hence, investigating resting-state brain activity, which is simple and task-free, is popular in studies aiming to develop biomarkers for distinguishing aMCI from healthy controls.^56,57^ However, recent studies found that task-induced functional connectivity may capture more behaviour-related information than resting-state functional connectivity even if the tasks during scanning are not directly relevant to the behavioral tasks.^58–60^ For example, Finn and Bandettini^59^ found that functional connectivity during movie watching could better predict watchers’ trait-like phenotypes in both cognition and emotion domains than resting-state connectivity. In the current study, we designed a paradigm for the passive perception of simple stimuli from three different modalities. Our MSEP paradigm is as simple, fast, and low in cognitive demand as the resting-state paradigm. Besides the well-performed classification, we also found robust correlations between sensory-induced functional connectivity and cognitive performance. These results support that sensory-induced neural activities could capture some changes in cognitive performance associated with aMCI, even though these cognitive functions are irrelevant to the perceptual stimuli per se.

Neural indices from all three modalities provide information for distinguishing between older adults with and without a diagnosis of aMCI. However, the amount of information varies between modalities, leading to different classification accuracies ranging from 69.6% to 92.4% (sensitivity: 79.5% to 95.5%; specificity: 57.1% to 88.6%). This is consistent with previous findings that neural responses to uni-modal stimuli could be used to classify aMCI from healthy controls.^41,61–63^ With our MSEP paradigm, we could obtain neural responses to different modalities in a single recording session, which enables us to combine informative component from different modalities for a better classification performance. Previous studies merging features from different modalities or sources usually combine features in the feature selection and model training stage and predict samples using the combined features. ^41,42,61^ The present study combines different features (i.e., the evoked response amplitude and functional connectivity) under three sensory conditions in the prediction output session. With our strategy, model prediction performance improved markedly in accuracy, sensitivity, and specificity (accuracy = 96.1%, sensitivity = 97.7%, specificity = 94.3%).

The first advantage of our pipeline is the high flexibility. In the model training and test stages, the SVM algorithm could be replaced with any other machine learning or deep learning algorithms, such as logistic regression or artificial neural networks. Then, permutation testing and the voting procedure could be performed in the same way as this study. Furthermore, the voting procedure is a highly flexible method to combine information from different features because it does not require merging different features before or during the model training and testing phases. Therefore, any model or pipeline that produces a prediction label for each sample will apply to the voting procedure. This feature provides the advantage that different neural indices could be trained and tested using different algorithms since the optimal algorithm is not necessarily the same across different neural indices due to their divergent data characteristics and task types.^64^

One of the main challenges in applying machine learning methods to psychological or medical research is interpretability.^65^ Another advantage of our analysis pipeline is providing insight into the neural mechanisms underlying successful classification. The adjusted CPM provides important edges that contribute to successful classification. Group comparison of neural indices among the correctly classified samples identifies significant components that distinguish the two groups. Therefore, these extracted components enable brain-behavior analyses to explore how changes in functional connectivity seen in aMCI are associated with cognitive impairment in this population.

Using PLV, we found that aMCI was associated with greater functional connectivity within central-frontal electrodes and greater functional connectivity from central-frontal to temporal electrodes, including parietal, temporal, and occipital electrodes in the theta to alpha frequency bands. Meanwhile, older adults diagnosed with aMCI exhibited greater whole-brain functional connectivity in the beta band. Our findings complement prior research on resting-state functional connectivity in MCI, showing hypo- and hyper-connectivity in MCI compared to controls.^56^ A recent meta-analysis of functional near-infrared spectroscopy studies showed that aMCI was associated with weaker functional connectivity between prefrontal and posterior parietal regions during resting state than healthy older adults.^66^ However, for task-induced functional connectivity, older adults with aMCI showed stronger connectivity between medial left frontal cortex and temporal regions compared to healthy controls during tasks of working memory and naming.^66^ These opposing results may reflect different constructs of brain functioning captured by resting-state functional connectivity and task-induced functional connectivity. Weaker functional connectivity at rest may suggest aMCI-related deficits in the stability of intrinsic organization. In comparison, higher functional connectivity may indicate a loss of neural complexity in aMCI in service of sensory perception and goal-oriented tasks. Our correlation analyses of the most important edges to cognitive tasks also support that the increased functional connectivity was correlated to impaired cognitive function in the aMCI group.

Besides functional connectivity, we also found that sensory-evoked response amplitude under all three sensory conditions are significant predictors of aMCI status. By comparing group differences in sensory-evoked responses, we found reduced amplitude at various latencies in all three sensory modalities. These results are consistent with previous studies investigating auditory- and visual-evoked response alterations in older adults diagnosed with MCI and early Alzheimer’s disease compared to healthy controls.^67–70^

Previous EEG studies found that older adults displayed enhanced early sensory-evoked response amplitude compared to young adults, which has been attributed to reflecting changes in sensory regulation mediated by the PFC.^17,36^ The decreased amplitude of sensory-evoked responses in individuals with aMCI may also reflect further deficits in frontal regulation of sensory responses attributed to early Alzheimer’s disease-related degeneration of frontal and temporal areas. This result is also consistent with deficits in frontal functioning in aMCI, such as inhibition,^2^ working memory,^71^ and executive function.^72^

This study introduces the novel MSEP paradigm suitable for the aMCI population and a classification analysis pipeline that enables researchers to combine features from different modalities flexibly and further explore the neural mechanism underlying the classification performance. Here, we employed the MSEP paradigm and classified older adults with aMCI from healthy older adults. Classification performance increased substantially after combining features from evoked potentials of multiple sensory modalities. Additionally, we found that the reduced sensory ERP amplitudes and increased functional connectivity in individuals with aMCI were the main contributors to the successful classification. These findings provide new insights into how EEG may be used to support the diagnosis and identification of neural biomarkers for aMCI and potentially other neurological disorders. Until now, past research has focused on examining sensory-perceptual processing in a single modality. Our study is the first to combine modalities to provide a more comprehensive picture of sensory-processing deficits, improving aMCI’s classification accuracy.

## Consent Statement

All participants provided informed consent for participation.

